# Genome-wide association study and genomic predictions for resistance against *Piscirickettsia salmonis* in coho salmon (*Oncorhynchus kisutch*) using ddRAD sequencing

**DOI:** 10.1101/124099

**Authors:** Agustín Barría, Kris A. Christensen, Katharina Correa, Ana Jedlicki, Jean P. Lhorente, William S. Davidson, José M. Yáñez

## Abstract

*Piscirickettsia salmonis* is one of the main infectious diseases affecting coho salmon (*Oncorhynchus kisutch*) farming. Current treatments have been ineffective for the control of the disease. Genetic improvement for *P. salmonis* resistance has been proposed as a feasible alternative for the control of this infectious disease in farmed fish. Genotyping by sequencing (GBS) strategies allow genotyping hundreds of individuals with thousands of single nucleotide polymorphisms (SNPs), which can be used to perform genome wide association studies (GWAS) and predict genetic values using genome-wide information. We used double-digest restriction-site associated DNA (ddRAD) sequencing to dissect the genetic architecture of resistance against *P. salmonis* in a farmed coho salmon population and identify molecular markers associated with the trait. We also evaluated genomic selection (GS) models in order to determine the potential to accelerate the genetic improvement of this trait by means of using genome-wide molecular information. 764 individuals from 33 full-sib families (17 highly resistant and 16 highly susceptible) which were experimentally challenged against *P. salmonis* were sequenced using ddRAD sequencing. A total of 4,174 SNP markers were identified in the population. These markers were used to perform a GWAS and testing genomic selection models. One SNP related with iron availability was genome-wide significantly associated with resistance to *P. salmonis* defined as day of death. Genomic selection models showed similar accuracies and predictive abilities than traditional pedigree-based best linear unbiased prediction (PBLUP) method.

## INTRODUCTION

Chile is the largest producer of coho salmon (*Oncorhynchus kisutch*) globally, reaching about 160,000 tons in 2014, representing more than 90% of total production (FAO 2016). However, the success and sustainability of this industry is constantly threatened by infectious diseases, including Salmon Rickettsial Syndrome (SRS). This disease is caused by *Piscirickettsia salmonis*, a gram-negative and facultative intracellular bacteria which was isolated for the first time in Chile in coho salmon (Cvitanich *et al*. 1991). Data from the Chilean National Fisheries and Aquaculture Service (Sernapesca) estimates that 59.3% of the morality rates ascribed to infectious disease were associated with SRS (Sernapesca 2016a). To date, control measures and treatments for SRS are based on antibiotics and vaccines. However, both strategies have not had the expected effectiveness under field conditions. Because of this, it is necessary to develop alternative strategies for the control of this disease (Yáñez *et al*. 2014a). In this regard, breeding for enhanced disease resistance is a feasible and sustainable option to improve animal health, welfare and productivity (Stear *et al*. 2001). A primary requisite for including disease resistance into a breeding program is the presence of significant additive genetic variation for the trait (Falconer and Mackay 1996). Commonly, data to evaluate resistance comes from experimental challenges carried out using siblings of the selection candidates (Ødegård *et al*. 2011;Yañez and Martinez 2010; Yáñez *et al*. 2014a). Quantitative studies have estimated significant genetic variation for resistance against different pathogens in salmonid species (Ødegård *et al*. 2011; Yáñez *et al*. 2014a). For instance, low to moderate heritabilities for resistance against *P. salmonis* in Atlantic salmon (*Salmo salar*) (h^2^ = 0.11 to 0.41) (Yáñez *et al*. 2013; Yáñez *et al*. 2014b) and coho salmon (h^2^ = 0.16) (Yañez *et al*. 2016a) have been estimated.

Marker assisted selection (MAS) can improve production traits in cases where the phenotypes are difficult to measure in the selection candidates (e.g. disease resistance traits) and the total additive genetic variance explained by genetic markers is high (Hayes and Goddard 2010). This methodology has been successfully applied for the improvement of resistance against infectious pancreatic necrosis in Atlantic salmon, which is controlled by a major quantitative trait locus (QTL) (Houston *et al*. 2012; Moen *et al*. 2015). In the case of polygenic traits, genomic selection (GS) (Meuwissen *et al*. 2001) can significantly improve selection accuracy of breeding values compared to traditional selection, and therefore enhance the response to selection for disease resistance in salmonid species (Tsai *et al*. 2016; Vallejo *et al*. 2016, 2017; Bangera *et al*. 2017; Correa *et al*. 2017).

Genotyping by sequencing (GBS) is an alternative for genotyping in cases when SNP panels are not available. This approach reduces the complexity of the genome, and can be used to identify thousands of markers without prior marker discovery efforts or a reference genome. Currently, several approaches of GBS have been developed, significantly reducing the cost and labor (Baird *et al*. 2008; Elshire *et al*. 2011; Peterson *et al*. 2012). These methodologies have been widely used in salmonid species, either to generate high density linkage maps (Brieuc *et al*. 2014; Gonen *et al*. 2014) or to perform association studies to identify genomic regions involved in the resistance against pathogens (Campbell *et al*. 2014; Palti *et al*. 2015b).

Double-digest restriction-site associated DNA (ddRAD) reduces DNA complexity by digesting DNA with two restriction enzymes (REs) simultaneously, and avoiding random shearing (Peterson *et al*. 2012). This approach has been widely used in genetic studies in aquaculture species (Robledo *et al*. 2017). In the present study, we used ddRAD sequencing to dissect the genetic architecture of resistance against *P. salmonis* in a farmed coho salmon population and identify molecular markers associated with the trait. Furthermore, GS models were used to evaluate the potential to accelerate the genetic improvement of resistance against *P. salmonis* in this coho salmon breeding population by means of using genome-wide molecular information.

## MATERIALS AND METHODS

### Coho Salmon Sampling

The coho salmon used in the present study were from the 2012 year-class of a genetic improvement program established in 1997 (owned by Pesquera Antares) and managed by Aquainnovo (Puerto Montt, Chile). Further details about this breeding population, in terms of reproductive management, rearing conditions, fish tagging, and breeding objectives are described by Yáñez *et al*. (2014c; 2016) and Dufflocq *et al*. (2016).

### Experimental Challenge

The experimental challenge is described in details by (Correa *et al*. 2015a) and Yáñez et al. (2016). Briefly, 2,607 individuals belonging to 108 maternal full-sib families (60 paternal half-sib families), were challenged against *Piscirickettsia salmonis*. Prior the challenge experiment, each fish was marked with a passive integrated transponder (PIT-tag), placed in the abdominal cavity for genealogy traceability during the challenge test. The experimental challenge test was performed at Aquainnovo’s Research Station, located in Lenca River, Xth Region, Chile. For the lethal dose 50 (LD_50_) calculation, a random sample of 80 fish were selected from the population. Four different dilutions from the *P. salmonis* inoculum were evaluated (1/10, 1/100, 1/1000 and 1/10000). Twenty fish per dilution were intraperitoneally (IP) injected with 0.2ml/fish. Daily mortality was recorded. This preliminary test spanned 26 days and a dilution of 1:680 was estimated as the LD_50_.

For the definitive challenge, fish were distributed into three tanks (7m^3^) with a salt water concentration of 31 ppt. An average of eight individuals (ranging from 1 to 18) from each of the 108 families were distributed into each tank.

The experimental challenge was performed through an intraperitoneal (IP) injection with 0.2ml/fish of the LD_50_ inoculum. The average weight of the fish at the inoculation was 279g (SD = 138g). Additionally, qRT-PCR was previously performed in order to control for the presence of Infectious Salmon Anemia Virus (ISAV), Infectious Pancreatic Necrosis Virus (IPNV) and *Flavobacterium spp*.

The challenge test was continued up to 50 days post injection. Throughout the challenge, environmental parameters (pH, temperature, salinity and oxygen) were measured and controlled. Daily mortality was removed from each tank, and a sample from anterior kidney was taken and stored at -80°C in RNALater. A necropsy assay was performed in conjunction with qRT-PCR to confirm the cause of death and the presence of *P. salmonis*. This was also done to control for the presence of other pathogens, such as *Vibrio ordalii, Renibacterium salmoninarum* and IPNV.

### Trait definitions and quantitative analysis

Resistance was defined as the day of death (DD) with values ranging from 1 to 50 depending on the time of death. Additionally, resistance was evaluated as a binary (BIN) trait, either dead or alive at the end of the challenge. Values for this trait were 1 in cases where the fish died during the challenge, or 0 if the fish survived until the end of the challenge. Initial Body Weight (IW) for each fish, was measured prior to the IP injection. A linear bivariate animal model was used in order to estimate covariance components for DD and BIN. Sex and tank were used as fixed effects, and IW was included as a covariate in the model. The animal model was fitted using ASREML (Gilmour *et al*. 2009). Heritability estimation was computed as follow:

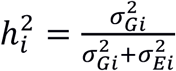
Where 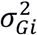 and 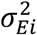 are the additive genetic and residual variances respectively.

The genetic correlation (*r_xy_*) between both traits was defined as Falconer and Mackay (1996):

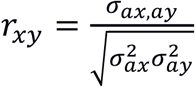
where *σ_ax,ay_* is the additive genetic covariance between traits, 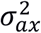 is the additive genetic variance of DD, and 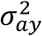 is the additive genetic variance of BIN.

### ddRAD library preparation and sequencing

Ten ddRAD libraries were produced by multiplexing 828 individuals following the protocol described by Peterson et al. (2012). For this, 64 parents (males and females) and 764 offspring representing the 17 most resistant and 16 most susceptible families were selected. An average of 23 (ranging from 11 to 43 individuals) offspring per family were chosen. Briefly, total DNA was extracted using the commercial kit Wizard SV Genomic DNA purification System (Promega) according to manufacturer’s protocol. Between 80 and 200 ng of DNA, from each individual was digested with two restriction enzymes (New England Biolabs, UK; NEB); 10 U of *Sbf*I (specific for the CCTGCA|GG motif)) and *Mse*I (specific for the T|TAA motif) in 12 μl reaction volume, including 1 μl of *Sbf*I and *Mse*I adapter (8.3 pM), for 90 min at 37°. The ligation reaction was carried out by adding 1 μl of T4 ligase (NEB) diluted 1:100 in T4 buffer and incubating for 150 min at 37° and subsequently at 16° overnight.

Each ligation mix was diluted with 189 μl of dilute TE buffer (1:10). Kodak DNA Polymerase (ABM), a high fidelity polymerase, was used to amplify DNA fragments with the correct adapters. PCR reactions (20 μl were prepared containing 10 μl of PCR mix 2x, 1 μl of primer mix (10 μM each), 6 μl of diluted ligation mix and 3 μl of nuclease-free water. Each sample was PCR amplified using the following conditions: 95° for 2 min, followed by 17 cycles of 95° for 20s, 66° for 30s and 68° for 40s. After PCR, amplicon quality was checked by loading 5 μl on a 2% agarose gel. Subsequently, samples were pooled, so that the final concentration was similar among them within each library. Each library was concentrated through an evaporation step for 80 min in a Centrivap Mobile Console Centrifugal Evaporator (Labconco). This step was conducted until 300 μl of the generated library was obtained. Final volume of each library was loaded on a 1% agarose gel. Size of the bands selected for sequencing ranged from 750 and 1500 bp and between 1,800 and 2,500 bp. DNA was purified through the QIAquick gel extraction kit (Qiagen) following manufacturer’s instructions. Finally, libraries were sequenced on an Illumina Hiseq2500 platform, using 150 base single-end sequencing.

### SNP identification

Raw sequence reads obtained from sequencing Illumina were analyzed using STACKS v. 1.41 (Catchen *et al*. 2011, 2013). This software was specifically developed to analyze short-read data generated through next generation sequencing (NGS) (Davey *et al*. 2013). Sample reads were trimmed to 134 bp for all subsequent analyses. Additionally, these reads were demultiplexed and filtered using *process_radtags*. Rad-tags which passed the quality filter were aligned to the *Oncorhynchus kisutch* reference genome (GenBank: MPKV00000000.1) using BWA v. 0.7.12 (Li and Durbin 2009). The reference genome was indexed and alignments were performed using the *mem* algorithm, all other parameters were set as default. Loci were then built using *pstacks* allowing a minimum depth of coverage of three to build a locus (-m 3). A catalog of loci was constructed through *cstacks* program using only the parents’ reads. To build the catalog, the maximum number of mismatches allowed between sample tags was three (-n 3), and the matching was based on genomic location (g). After this, the *sstacks* program was used in order to match rad-tags against the catalog based again on genomic location (g), followed by *populations* software, using defaults parameters. Loci were considered as valid if they are present, in at least, 75% of the individuals of the population.

### Genomic Association Study

In order to associate the molecular markers to *P. salmonis* resistance, either as DD or BIN, a GWAS was performed using the GenABEL library (Aulchenko *et al*. 2007) implemented in R (http://www.r-project.org). Prior to analyses, identified SNPs were submitted to a Quality Control (QC), with the following parameters: Minor Allele Frequency (MAF) ≥ 0.01, Hardy-Weinberg Equilibrium (HWE) = p < 1×10^−6^, individual call rate > 0.70 and SNP call rate > 0.90. Additionally, all markers were used in order to estimate identity by state (IBS). Association analysis was performed through the Grammar-Gamma method (Svishcheva *et al*. 2012) including a genomic kinship matrix estimated using filtered SNPs. The polygenic function (Thompson and Shaw 1990) was used to fit two different univariate additive polygenic models. The linear regression model was defined as follows:

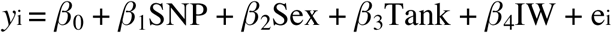
Where *y_i_* is the vector of phenotypes (DD), *β*_0_ is the mean, *β*_1_ is the effect of each SNP, *β*_2_ is the fixed effect of the sex, *β*_3_ is the fixed effect of the tank, *β*_4_ is the effect of the initial weight as covariate and e_i_ is the random residual.

The logistic regression models for BIN was:

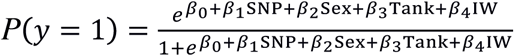
Where *P* is the probability of the random variable to be one, *β*_0_ is the mean, *β*_1_ is the effect of each SNP, *β*_2_ is the fixed effect of the sex, *β*_3_ is the fixed effect of the tank, *β*_4_ is the effect of the initial weight as covariate and e_i_ is the random residual.

Genomic-wide significance was assessed by False Discovery Rate (FDR) (Benjamini and Hochberg 1995). Briefly, *p* values were ordered from p_1_ ≤ p_2_ ≤… ≤ p_k_ where *k* is the number of SNPs tested. Beginning from the largest *p* value, the first p value (p_i_) that satisfied: p_i_ ≤ /*k*\*0.05, where i = ith observation. Finally, the p_i_ that satisfied the condition became the significant value.

Finally, *BLASTn* was used to align the genome-wide associated region, against Atlantic salmon genome (Genbank: GCA_000233375.4) and identify candidate genes associated with *P. salmonis* resistance.

### Genomic Prediction

The pedigree-based approach, PBLUP was used as the control for the genomic evaluations, and EBV for each individual were estimated using a linear mixed model implemented in BLUPF90 (Misztal *et al*. 2016). The model used was as follows:

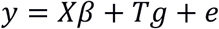
where *y* is a vector of phenotypes (BIN or DD), *β* is a vector of fixed effects (Sex, Tank and initial weight effects), *g* is a vector of random additive polygenic genetic effects that follows a normal distribution 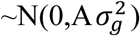, *X* and *T* are incidence matrices, *A* is the additive relationship matrix, and *e* is the residual (Lynch and Walsh 1998).

The genomic EBV (GEBV) for each individual were estimated using GBLUP (*Genomic* BLUP) as implemented in the BLUPF90 software. GBLUP is a modification of the PBLUP method, where the numerator relationship matrix *A* is replaced by a genomic relationship matrix *G*, as described by VanRaden (2008). Pedigree (PBLUP) and genomic (GBLUP) heritabilities were calculated using the AIREMLF90 software (Misztal *et al*. 2016) as previously described.

The different models were compared using a five-fold cross validation scheme. Briefly, all genotyped and phenotyped animals were randomly separated into five validations sets, which were predicted one at a time by masking their phenotypes and using the remaining animals as a training set to estimate the marker effects. Thus, for each validation run, the dataset was split into a training set (80%) and a validation set (20%). To reduce the stochastic effects, this cross-validation analysis was replicated 10 times. Accuracy was used to assess the performance of each model and was estimated as:

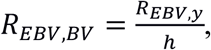
where *R_EBV,y_* is the correlation between the EBV of a given model (predicted for the validation set using information from the training set) and the actual phenotype, while *h* is the square root of the pedigree-based estimate of heritability (Legarra *et al*. 2008; Ødegård *et al*. 2014). To test prediction accuracies obtained using different SNP densities, 1 K, 2 K, 3 K and 4 K SNPs were randomly selected from the SNPs that passed the quality control. Finally, accuracies were calculated for each model and SNP density, and compared to those obtained with the PBLUP model.

### Data availability

Table S1 contains genotypic data. Data of pedigree and phenotypes that supports the findings in this study are available from Aquainnovo and AquaChile but restrictions apply to the availability of these data, which were used under license for the current study, and so are not publicly available. However, data are available from the corresponding author upon reasonable request and with permission of Aquainnovo and AquaChile.

## RESULTS

### Challenge test

Mortality began on day 10 post challenge, with evident clinical signs and pathological lesions typical of the Salmon Rickettsial Syndrome. These signs include swollen kidney, splenomegaly and liver with a yellowish tone (Rozas and Enríquez 2014). Challenged families showed considerable phenotypic variation for *P. salmonis* resistance. Average mortality of all families reached 41% during the 50-day challenge. Average cumulative mortality rate among the 17 best and 16 worst families selected for genotyping, reached 19% and 63%, respectively (Figure 1).

**Figure 1.**
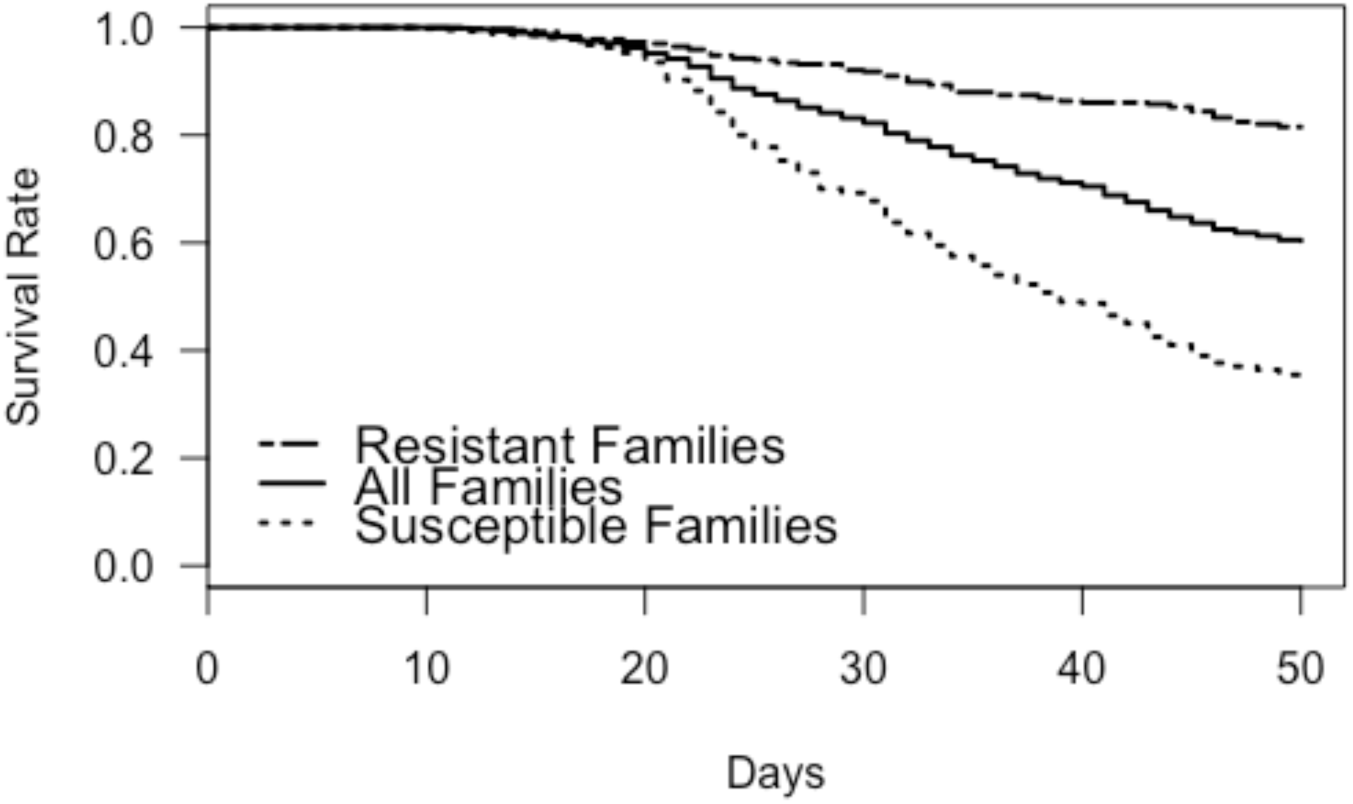
Kaplan-Meier curves for *Pisciricketssia salmonis* experimental challenge in coho salmon. Average mortality curves for the 108 full-sib families, and the 17 best and 16 worst families.

### Heritabilities and correlations between traits

Moderate significant additive genetic variation was estimated for both DD and BIN. Using a pedigree relationship matrix, h^2^ was estimated as 0.26 (± 0.01) and 0.42 (± 0.01) for DD and BIN, respectively. In the case of using a G matrix, the h^2^ was estimated to be 0.32 and 0.38 for DD and BIN, respectively. Genetic correlation between traits was very high and significantly different from zero (-0.95 ± 0.03). Additionally, high significant phenotypic correlation was estimated, being -0.77 (± 0.01).

### ddRAD sequencing

Prior to quality control (QC), per base quality (Phred score) was evaluated. The average quality score ranged from 36 to 38 among libraries, indicating high quality of data. Illumina sequencing, including parents, yielded an average of 156,058,078 (± 16 millions) of raw sequences. After QC, which included the removal of low quality sequences and reads with either missing or ambiguous barcodes, an average of 31,660,024 of the reads were removed. In parallel with QC, reads were trimmed to 134 bp. In this regard, 79% of the raw reads were retained for subsequent analysis. In order to create a set of all possible alleles in the population, data sets of the parental samples were used to create the STACKS catalogue. This catalogue consisted of 106,309 unique ddRAD loci from which 55,770 markers from 757 individuals were obtained.

### Association Analysis

A total of 4,174 markers and 592 animals passed all QC criteria (see Table S1). Association analysis identified one marker showing genome-wide statistical significance (p value = 5.50E-05) (Fig. 2) for DD. This marker was located on scaffold01025. However, none of the markers were significant using the FDR significance threshold when resistance was defined as a binary trait (Fig. 3). For this trait only one marker was identified as suggestively associated (p value = 1.50E-05). This marker was located in Chromosome 29. BLAST results determined that the genomic region surrounding the significant SNP associated with DD, was located on chromosome Ssa03, proximate to a *heme oxygenase 2-like* gen. Location, proportion of heritability and phenotypic variance explained, and p-values for the significant and suggestive marker are summarized in table 1.

**Figure 2.**
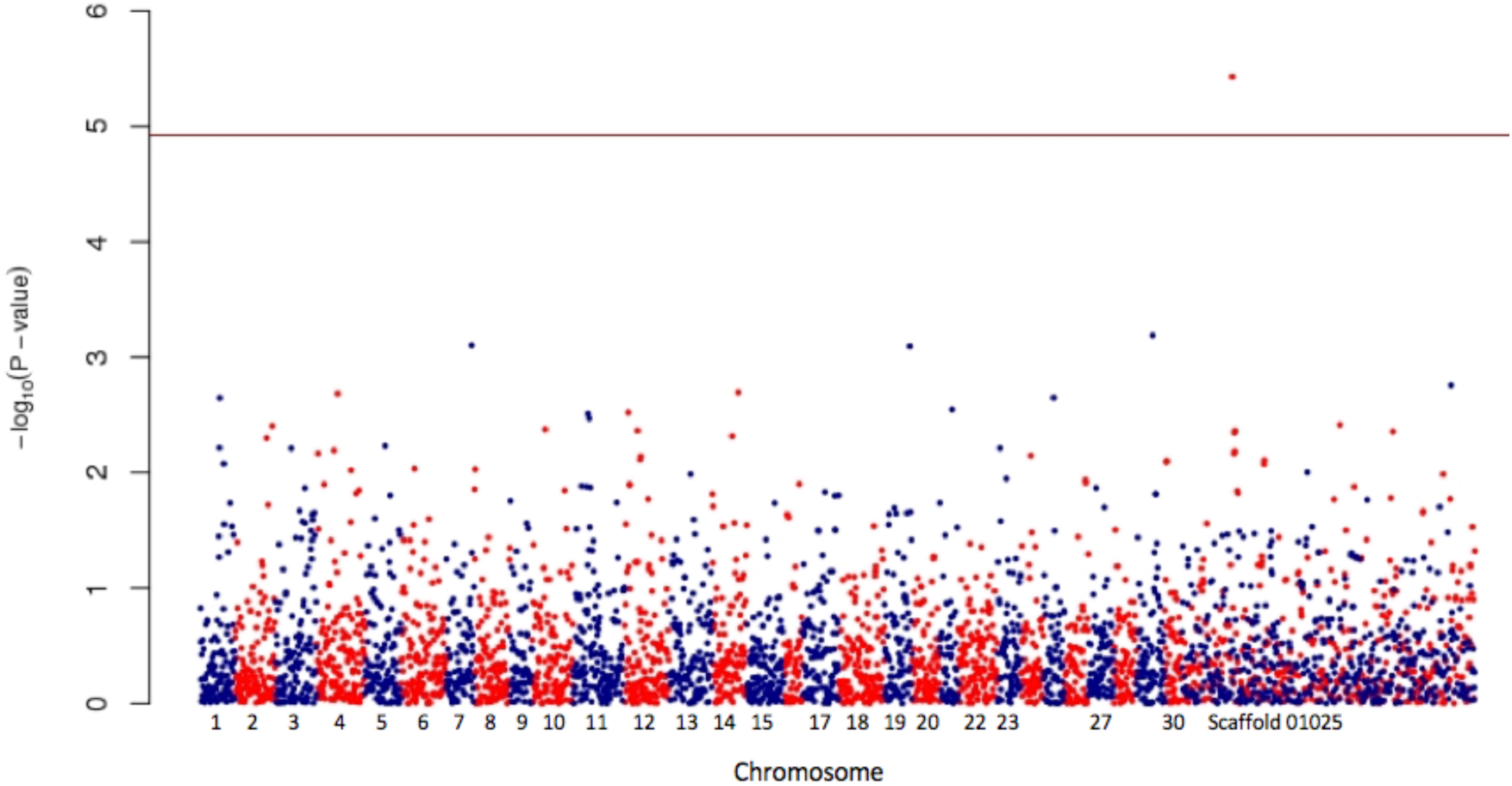
Genomic association analysis for resistance against *Pisciricketssia salmonis* in coho salmon population. Resistance was defined as day of death. The horizontal line indicates the FDR significance threshold.

**Figure 3.**
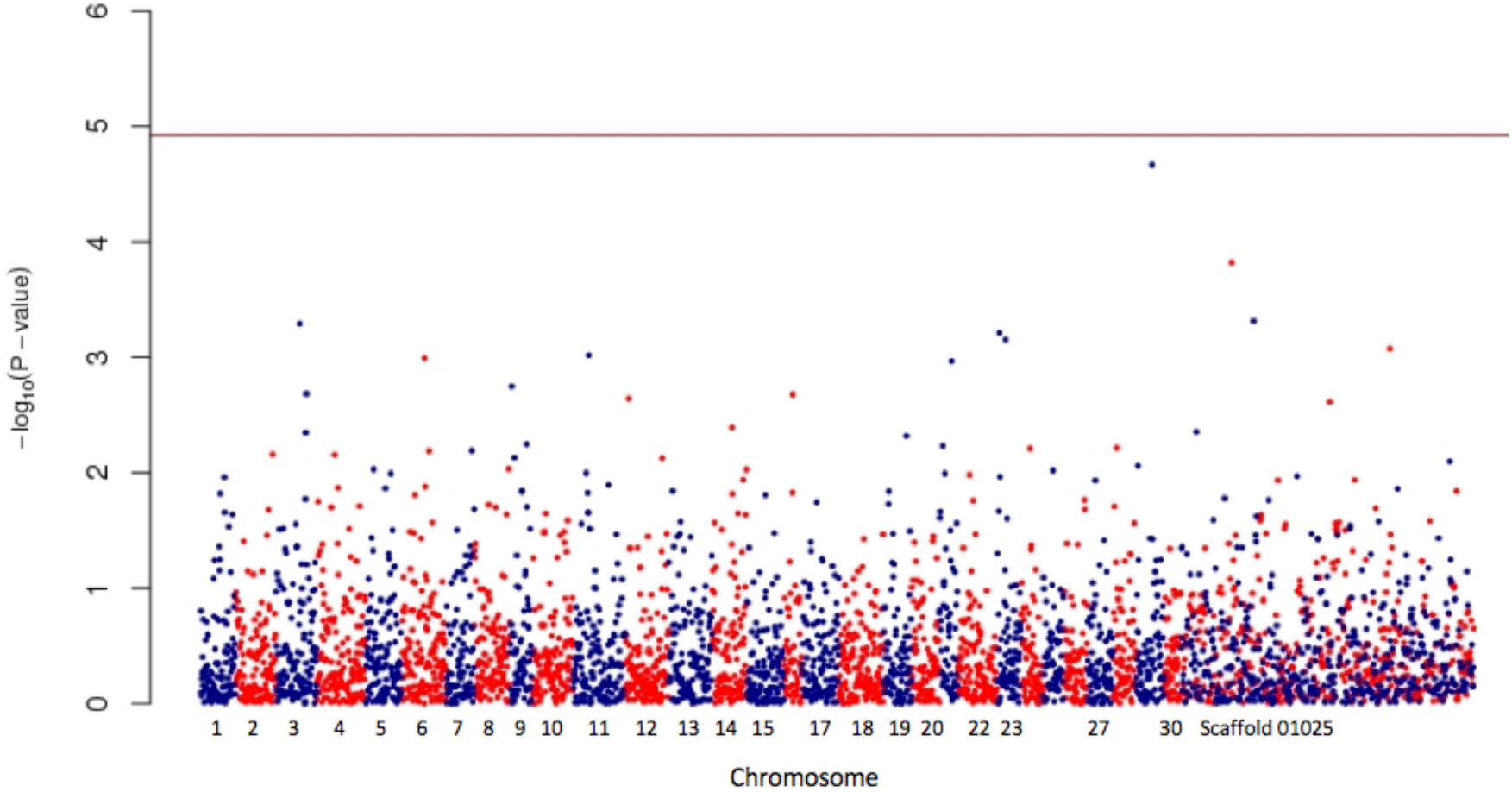
Genomic association analysis for resistance against *Pisciricketssia salmonis* in coho salmon population. Resistance was defined as a binary trait. The horizontal line indicates the FDR significance threshold.

**Table 1.**
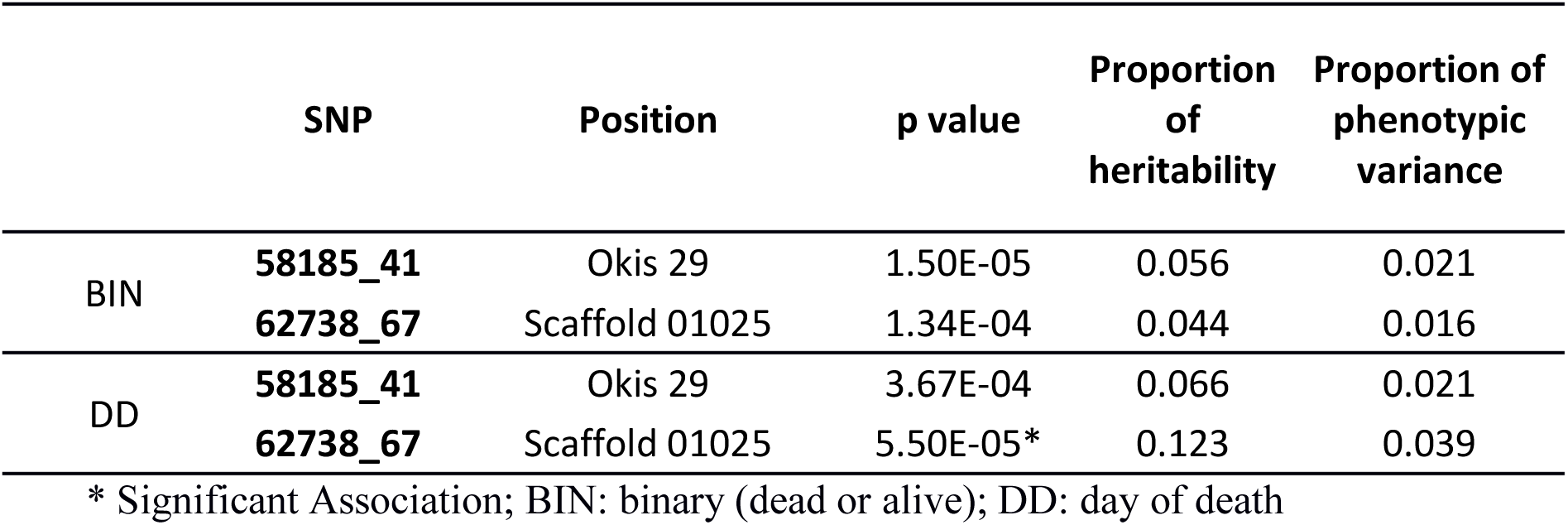
Location, p value, proportion of heritability and phenotypic variance explained by the two most important markers associated with resistance against *P. salmonis*.

### Genomic Prediction

Prediction was assessed using phenotype and genotype information from 4,174 SNPs which passed all QC steps from 592 individuals. These individuals were randomly split into five different training (474) and validation (118) sets, either for DD or BIN. PBLUP accuracy reached 0.58 and 0.61 for DD and BIN, respectively. These values were similar to those obtained using genomic information, ranging from 0.57-0.60 for DD and 0.60-0.62 for BIN at different SNPs densities (Table 2). Moreover, using only phenotypic information, reliability was also higher for BIN, reaching 0.38 in contrast to 0.36 for DD. When genomic data was used, reliability values were similar to those obtained with PBLUP ranging from 0.34 to 0.39. Finally, predictive ability reached up to 38% in the case of DD, showing an increase around 15% in the case of BIN, independently of the use of genotype information.

**Table 2.**
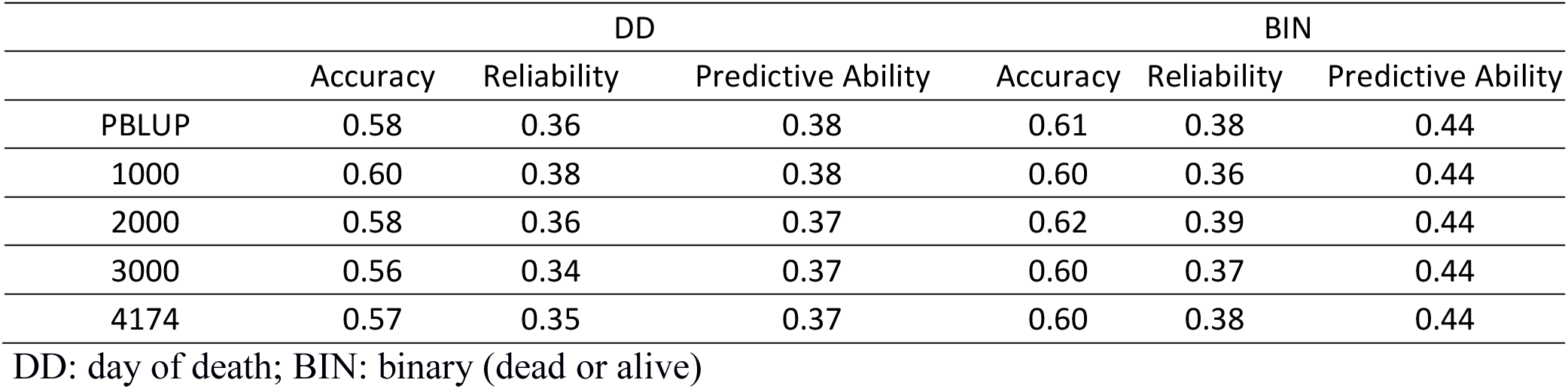
Accuracy, reliability and predictive ability of GEBVs assessed by five-fold cross validation for resistance against *Piscirickettsia salmonis* defined as day of death or as binary trait in coho salmon population using four SNPs densities.

## DISCUSSION

*Piscirickettsia salmonis* is the etiological agent of SRS, the main disease affecting salmon production in Chile. Currently, control strategies have not shown the expected effectiveness in field conditions (Rozas and Enríquez 2014). Some strains of *P. salmonis* have presented resistance to Quinolones and oxytetracycline and intermediate susceptibility to florfenicol (Henríquez *et al*. 2016), which are some of the most used antibiotics in the Chilean industry for the treatment of this infectious disease (nearly 90% of the total amount) (Sernapesca 2016b).

Selective breeding has been successfully applied commercially for important production traits of aquaculture species, including salmonids (Gjedrem 2012; Yáñez *et al*. 2014a). Genetic improvement has included disease resistance, based on information of relatives, but does not allow for the use of within-family genetic variation and consequently affects the achievable genetic progress due to the lower accuracy on EBVs (Falconer and Mackay 1996). GS has had a major impact on traits difficult to measure on the selection candidates themselves, as is the case of disease resistance, and has improved genetic gain and selection accuracy (Goddard and Hayes 2009). In this sense, breeding schemes can incorporate genomic information, either through GS or via MAS, in order to accelerate the genetic progress for disease resistance traits (Yáñez *et al*. 2015).

Significant genetic variation for *P. salmonis* resistance was detected in the present study. Moderate heritabilities were estimated using different trait definitions, either for DD or BIN using both A and G matrices. Estimated heritabilities were higher for resistance as binary trait when compared to DD, independent of the matrix used in the analysis. Additionally, high genetic correlation between traits was estimated (-0.95 ± 0.03), suggesting that DD and BIN, are likely measures of the same trait.

Previously, Yáñez (2016), estimated a heritability of 0.16 for resistance against *P. salmonis*, defined as day of death, in the same coho salmon population, through a linear model. However, our estimations used a directed sample of the population, which explains the difference between estimations. These results are in agreement with previous findings which estimated heritabilities for resistance against *P. salmonis* in Atlantic salmon ranging from 0.11 to 0.41 (Yáñez *et al*. 2013, 2014b; Correa *et al*. 2015b). The genetic variation for *P. salmonis* resistance, and values of heritability, are in accordance with different studies that also found significant genetic resistance against other bacterial disease in salmonid species (Gjøen *et al*. 1997; Ødegård *et al*. 2006; Vallejo *et al*. 2016).

Association analysis identified one molecular marker significantly associated with *P. salmonis* resistance defined as day of death. The reference genome of coho salmon is not completely assembled into chromosomes (∼75% anchored), and it was not possible to elucidate to which chromosome this marker belongs. However, the availability of a high quality *Salmo salar* genome reference (Lien *et al*. 2016) made it possible to identify the marker on a genomic region of chromosome Ssa03 proximate to *Hem Oxygenase-2* (HO2), an enzyme involved in the iron accessibility through the degradation of a heme group (Kikuchi *et al*. 2005). Different studies have already demonstrated the ability of *P. salmonis* to evade the immune system and infect, survive and replicate inside macrophages (McCarthy *et al*. 2008; Rojas *et al*. 2009). Transcriptional studies performed by Rise *et al*. (2004) showed that after *P. salmonis* infection, ferritin and transferrin, both related with binding and the storing of iron, were up regulated in Atlantic salmon macrophages. Similar results were obtained by Pulgar *et al*. (2015) who showed that after an IP challenge against *P. salmonis*, genes related to iron homeostasis were differentially up-regulated in head kidney of resistant families of Atlantic salmon compared with the susceptible ones. Moreover, head-kidney iron content was significantly lower in resistant families, which was positive correlated with *P. salmonis* load. This evidence makes it possible to suggest that the deprivation of the intracellular iron content is a key immune response mechanism against *P. salmonis* infection, and may reduce its replication rate. A similar mechanism was previously demonstrated to reduce bacterial growth inside murine macrophages for a variety of bacterial infections (Paradkar *et al*. 2011). *Hem oxygenase 1* has been shown to be up-regulated 2 hr after an infection generated by the gram negative bacteria, *Salmonella typhimurium* (Nairz *et al*. 2007). This gene was able to reduce oxidative stress, has been involved in murine-macrophages iron export and a subsequent intracellular iron deprivation (making iron inaccessible to bacteria) (Nairz *et al*. 2007). These results support the potential role that *HO2* could be playing in the resistance against *P. salmonis* in coho salmon (i.e. limiting bacteria replication rate through iron deprivation at the intra-cellular level). However, further studies need to be performed in order to have a better understanding of the host immune response against the infection and confirm the role of *HO2* in the variation of *P. salmonis* resistance.

Previously, resistance against bacterial diseases has been suggested as a polygenic trait in aquaculture species. For example, Palaiokostas (2016) suggested resistance against *Photobacterium damselae subsp*. had a polygenic genetic architecture in Gilthead Sea Bream (*Sparus aurata*). Using a 50K SNP genotyping array, it was possible to elucidate a moderately polygenic architecture for resistance against *P. salmonis* in Atlantic salmon (Correa *et al*. 2015b). In the current study, and using 4 K SNPs, a moderate polygenic genetic architecture for resistance against *P. salmonis* in coho salmon population is suggested. However, due to the low marker density, it might be possible that this density is not enough to capture linkage disequilibrium (LD) between genetic markers and all the important QTLs controlling *P. salmonis* resistance in coho salmon.

Currently, GS has been demonstrated to be a feasible alternative over traditional breeding methods to increase response to selection due to higher accuracy breeding values, in both, simulated and empirical studies in aquaculture species (Sonesson and Meuwissen 2009; Tsai *et al*. 2015, 2016, Vallejo *et al*. 2016, 2017; Bangera *et al*. 2017; Correa *et al*. 2017).

The current study, showed similar accuracies, reliabilities and predictive ability (PA) when comparing PBLUP and GBLUP (even among different SNPs densities). In this regard, estimated accuracies with PBLUP were 0.58 and 0.60, which are highly similar to those obtained using genomic data, from 0.56 to 0.62 (independent of the number of genetic markers used). Similar tendencies were obtained for reliability and PA, where the average values were identical between PBLUP and GBLUP.

When resistance was defined as a binary trait, both accuracy and reliability, were slightly higher in comparison with resistance as day of death. However, PA was almost 15% higher for BIN than DD, either with or without genomic data, reaching up to 0.44. These results, are similar to those obtained in GS for Bacterial Cold Water Disease (BCWD), in rainbow trout (*Oncorhynchus mykiss*), using SNP arrays and RAD seq (Vallejo *et al*. 2016), obtaining similar PA values with or without genotypic information, ranging from 0.37-0.50 and from 0.26-0.41 for DD or BIN, respectively. Vallejo *et al*. (2016) attributed this results to the small training sample size of their study (n=583), which is higher than that used in the current study (n=473). In this regard, PA increase reached up to 108% relative to PBLUP when the number of phenotyped and genotyped individuals increase, even when less SNPs were used (Vallejo *et al*. 2017). Interestingly, estimated PA for BCWD resistance are similar using 10 K SNPs obtained by RAD as those obtained using a 40 K SNP array, even if resistance is defined as day of death or as binary. Authors attributed this to the long-range LD within the population (Vallejo *et al*. 2016).

In case of Atlantic salmon, improvement in accuracy raised 20% using 112 K SNPs either for length and weight in juvenile individuals (Tsai *et al*. 2015). Moreover, sea lice resistance has showed a relative improvement in accuracy of 22 and 27%, relative to PBLUP, using 37 K and 33 K SNPs respectively. Additionally, improvement in reliability reached up to 52% with 220 K (Ødegård *et al*. 2014; Tsai *et al*. 2016; Correa *et al*. 2017). In case of *P. salmonis* resistance, relative reliability has been improved in 25 and 30% either for resistance, defined as day of death or as a binary trait respectively (Bangera *et al*. 2017).

Using 2b-RAD methodology, Palaiokostas (2016) showed a reduction in the improvement of accuracies, for resistance against a bacterial disease in *S. aurata*, when lower SNPs densities were evaluated. In this regard, an improvement of 53% in the estimated accuracy was observed using 12 K molecular markers, raising it up to 0.46 in comparison with 0.30 obtained through PBLUP through a Bayesian model. However, using 2 K SNPs, accuracy showed a maximum increase of 20%. However, this improvement was completely abolished when the number of markers was reduced to 700.

We hypothesize that the similar results with or without genomic information in the current study, could be due to the small training sample size and the low density of SNPs markers used, making it difficult to capture the LD between markers and all of the loci influencing this trait. Moreover, simulations performed by Perez Enciso (2015), suggest that even using a higher number of molecular markers obtained by RAD seq, accuracies are lower than using a medium density SNP array, likely due to that in the RAD seq methodology; SNPs are tightly linked among them, whereas in arrays SNPs are uniformly distributed along the genome. The availability of dense SNP arrays for coho salmon, as it is already the case for Atlantic salmon (Houston *et al*. 2014; Yañez *et al*. 2016b) and rainbow trout (Palti *et al*. 2015a), may allow increase the accuracy for predicting genomic breeding values and the power for the determination of the genetic factors involved in economically-important traits, including *P. salmonis* resistance, in this species.

## 5. Conclusions

Moderate significant genetic variation was estimated for resistance against *Piscirickettsia salmonis* in coho salmon, using either pedigree or genomic information. These results highlight the feasibility to include this character into genetic improvement programs. One SNP was genome-wide significantly associated with resistance to *P. salmonis*, and may be involved in the immune host response against this infection through iron homeostasis mechanism. Genomic predictions using ddRAD genotypes including 4,174 loci showed similar accuracies as PBLUP. To our knowledge, this is the first study aiming at dissecting the genetic architecture of resistance against *P. salmonis*, in coho salmon population.

## Acknowledgements

AB and KC want to acknowledge the National Commission of Scientific and Technologic Research (CONICYT) for the funding through the National PhD funding program. AB wants to acknowledge the Government of Canada for the funding through the Canada-Chile Leadership Exchange Scholarship. This project was funded by the U-Inicia grant, from the Vicerrectoria de Investigación y Desarrollo, Universidad de Chile. This work has been conceived on the frame of the grant FONDEF NEWTON-PICARTE (IT14I10100), funded by CONICYT (Government of Chile) and the Newton Fund - The British Council (Government of United Kingdom).

## Authors’ contributions

AB performed DNA extraction, library construction, ddRAD analysis, GWAS analysis and wrote the initial version of the manuscript. KrC performed library construction and contributed on the data analysis. KC performed Genomic prediction analysis. AJ performed DNA extraction. JPL contributed with study design. WD contributed with analysis and discussion. JMY conceived and designed the study, supervised work of AB and contributed to the analysis, discussion and writing. All authors have reviewed and approved the manuscript.

## Bibliography

Aulchenko, Y. S., S. Ripke, A. Isaacs, and C. M. van Duijn, 2007 GenABEL: An R library for genome-wide association analysis. Bioinformatics 23: 1294–1296.

Baird, N. A., P. D. Etter, T. S. Atwood, M. C. Currey, A. L. Shiver et al., 2008 Rapid SNP discovery and genetic mapping using sequenced RAD markers. PLoS One 3: 1–7.

Bangera, R., K. Correa, J. P. Lhorente, R. Figueroa, and J. M. Yáñez, 2017 Genomic predictions can accelerate selection for resistance against *Piscirickettsia salmonis* in Atlantic salmon (*Salmo salar*). BMC Genomics 18: 121.

Benjamini, Y., and Y. Hochberg, 1995 Controlling the False Discovery Rate: A Practical and Powerful Approach to Multiple Testing. J. R. Stat Soc. 57: 289–300.

Brieuc, M. S. O., C. D. Waters, J. E. Seeb, and K. a Naish, 2014 A dense linkage map for Chinook salmon (*Oncorhynchus tshawytscha*) reveals variable chromosomal divergence after an ancestral whole genome duplication event. G3 (Bethesda). 4: 447–460.

Campbell, N. R., S. E. LaPatra, K. Overturf, R. Towner, and S. R. Narum, 2014 Association mapping of disease resistance traits in rainbow trout using restriction site associated DNA sequencing. G3 (Bethesda). 4: 2473–81.

Catchen, J. M., A. Amores, P. Hohenlohe, W. Cresko, J. H. Postlethwait et al., 2011 Stacks: Building and Genotyping Loci De Novo From Short-Read Sequences. G3 (Bethesda). 1: 171–182.

Catchen, J., P. A. Hohenlohe, S. Bassham, A. Amores, and W. A. Cresko, 2013 Stacks: An analysis tool set for population genomics. Mol. Ecol. 22: 3124–3140.

Correa, K., R. Bangera, R. Figueroa, J. P. Lhorente, and J. M. Yáñez, 2017 The use of genomic information increases the accuracy of breeding value predictions for sea louse (*Caligus rogercresseyi*) resistance in Atlantic salmon (*Salmo salar*). Genet. Sel. Evol. 49: 15.

Correa, K., M. Filp, D. Cisterna, M. E. Cabrejos, C. Gallardo-Escárate et al., 2015a Effect of triploidy in the expression of immune-related genes in coho salmon *Oncorhynchus kisutch* (Walbaum) infected with *Piscirickettsia salmonis*. Aquac. Res. 46: 59–63.

Correa, K., J. Lhorente, M. Lopez, L. Bassini, S. Naswa et al., 2015b Genome-wide association analysis reveals loci associated with resistance against *Piscirickettsia salmonis* in two Atlantic salmon (*Salmo salar L*.) chromosomes. BMC Genomics 16: 854.

Cvitanich, J., O. Garate, and C. E. Smith, 1991 The isolation of a rickettsia-like organism causing disease and mortality in Chilean salmonids and its confirmation by Koch’s postulate. J. Fish Dis. 14: 121–146.

Davey, J. W., T. Cezard, P. Fuentes-Utrilla, C. Eland, K. Gharbi et al., 2013 Special features of RAD Sequencing data: Implications for genotyping. Mol. Ecol. 22: 3151–3164.

Dufflocq, P., J. P. Lhorente, R. Bangera, R. Neira, S. Newman et al., 2016 Correlated response of flesh color to selection for harvest weight in coho salmon (*Oncorhynchus kisutch*). Aquaculture 6–11.

Elshire, R. J., J. C. Glaubitz, Q. Sun, J. A. Poland, K. Kawamoto et al., 2011 A robust, simple genotyping-by-sequencing (GBS) approach for high diversity species. PLoS One 6: 1–10.

Falconer, D. S., and T. F. C. Mackay, 1996 Quantitative Genetics.

FAO, 2016 Fisheries and Aquaculture Information and Statistical Branch.

Gilmour, A. R., B. J. Gogel, B. R. Cullis, and R. Thompson, 2009 ASReml User Guide release 3.0. 372 p. VSN Int. Ltd, Hemel Hempstead, UK 372.

Gjedrem, T., 2012 Genetic improvement for the development of efficient global aquaculture: A personal opinion review. Aquaculture 344–349: 12–22.

Gjøen, H. M., T. Refstie, O. Ulla, and B. Gjerde, 1997 Genetic correlations between survival of Atlantic salmon in challenge and field tests. Aquaculture 158: 277–288.

Goddard, M., and B. Hayes, 2009 Mapping genes for complex traits in domestic animals and their use in breeding programmes. Nat. Rev. Genet. 10: 381–391.

Gonen, S., N. R. Lowe, T. Cezard, K. Gharbi, S. C. Bishop et al., 2014 Linkage maps of the Atlantic salmon (*Salmo salar*) genome derived from RAD sequencing. BMC Genomics 15: 166.

Hayes, B., and M. Goddard, 2010 Genome-wide association and genomic selection in animal breeding. Genome 876–883.

Henríquez, P., M. Kaiser, H. Bohle, P. Bustos, and M. Mancilla, 2016 Comprehensive antibiotic susceptibility profiling of Chilean *Piscirickettsia salmonis* field isolates. J. Fish Dis. 39: 441–448.

Houston, R. D., J. W. Davey, S. C. Bishop, N. R. Lowe, J. C. Mota-Velasco et al., 2012 Characterisation of QTL-linked and genome-wide restriction site-associated DNA (RAD) markers in farmed Atlantic salmon. BMC Genomics 13: 244.

Houston, R. D., J. B. Taggart, T. Cézard, M. Bekaert, N. R. Lowe et al., 2014 Development and validation of a high density SNP genotyping array for Atlantic salmon (*Salmo salar*). BMC Genomics 15: 90.

Kikuchi, G., T. Yoshida, and M. Noguchi, 2005 Heme oxygenase and heme degradation. Biochem. Biophys. Res. Commun. 338: 558–567.

Legarra, A., C. Robert-Granié, E. Manfredi, and J. M. Elsen, 2008 Performance of genomic selection in mice. Genetics 180: 611–618.

Li, H., and R. Durbin, 2009 Fast and accurate short read alignment with Burrows-Wheeler transform. Bioinformatics 25: 1754–1760.

Lien, S., B. F. Koop, S. R. Sandve, J. R. Miller, P. Matthew et al., 2016 The Atlantic salmon genome provides insights into rediploidization. Nature 533: 200–205.

Lynch, M., and B. Walsh, 1998 Genetics and analysis of quantiative traits. Sunderland: Sinauer Associates.

McCarthy, Ú. M., J. E. Bron, L. Brown, F. Pourahmad, I. R. Bricknell et al., 2008 Survival and replication of *Piscirickettsia salmonis* in rainbow trout head kidney macrophages. Fish Shellfish Immunol. 25: 477–484.

Meuwissen, T. H., B. J. Hayes, and M. E. Goddard, 2001 Prediction of Total Genetic Value Using Genome-Wide Dense Marker Maps. Genetics 157: 1819–1829.

Misztal, I., S. Tsuruta, D. Lourenco, Y. Masuda, I. Aguilar et al., 2016 Manual for BLUPF90 family of programs.

Moen, T., J. Torgersen, N. Santi, W. S. Davidson, M. Baranski et al., 2015 Epithelial Cadherin Determines Resistance to Infectious Pancreatic Necrosis Virus in Atlantic Salmon. Genetics 200: 1313–26.

Nairz, M., I. Theurl, S. Ludwiczek, M. Theurl, S. M. Mair et al., 2007 The co-ordinated regulation of iron homeostasis in murine macrophages limits the availability of iron for intracellular *Salmonella typhimurium*. Cell. Microbiol. 9: 2126–2140.

Ødegård, J., M. Baranski, B. Gjerde, and T. Gjedrem, 2011 Methodology for genetic evaluation of disease resistance in aquaculture species: Challenges and future prospects. Aquac. Res. 42: 103–114.

Ødegård, J., T. Moen, N. Santi, S. A. Korsvoll, S. Kjøglum et al., 2014 Genomic prediction in an admixed population of Atlantic salmon (*Salmo salar*). Front. Genet. 5: 1–8.

Ødegård, J., I. Olesen, B. Gjerde, and G. Klemetsdal, 2006 Evaluation of statistical models for genetic analysis of challenge test data on furunculosis resistance in Atlantic salmon (*Salmo salar*): Prediction of field survival. Aquaculture 259: 116–123.

Palaiokostas, C., S. Ferarreso, R. Franch, R. D. Houston, and L. Bargelloni, 2016 Genomic prediction of resistance to pasteurellosis in gilthead sea bream (*Sparus aurata*) using 2b-RAD sequencing. G3 (Bethesda). X: 1–8.

Palti, Y., G. Gao, S. Liu, M. P. Kent, S. Lien et al., 2015a The development and characterization of a 57K single nucleotide polymorphism array for rainbow trout. Mol. Ecol. Resour. 15: 662–672.

Palti, Y., R. L. Vallejo, G. Gao, S. Liu, A. G. Hernandez et al., 2015b Detection and Validation of QTL Affecting Bacterial Cold Water Disease Resistance in Rainbow Trout Using Restriction-Site Associated DNA Sequencing. PLoS One 10: e0138435.

Paradkar, P. N., I. De Domenico, N. Durchfort, I. Zohn, J. Kaplan et al., 2011 Iron depletion limits intracellular bacterial growth in macrophages Iron depletion limits intracellular bacterial growth in macrophages. Blood 112: 866–874.

Peterson, B. K., J. N. Weber, E. H. Kay, H. S. Fisher, and H. E. Hoekstra, 2012 Double digest RADseq: An inexpensive method for de novo SNP discovery and genotyping in model and non-model species. PLoS One 7:.

Pulgar, R., C. Hödar, D. Travisany, A. Zuñiga, C. Domínguez et al., 2015 Transcriptional response of Atlantic salmon families to Piscirickettsia salmonis infection highlights the relevance of the iron-deprivation defence system. BMC Genomics 16: 495.

Rise, M. L., S. R. M. Jones, G. D. Brown, K. R. von Schalburg, W. S. Davidson et al., 2004 Microarray analyses identify molecular biomarkers of Atlantic salmon macrophage and hematopoietic kidney response to *Piscirickettsia salmonis* infection. Physiol. Genomics 20: 21–35.

Robledo, D., C. Palaiokostas, L. Bargelloni, P. Martínez, and R. Houston, 2017 Applications of genotyping by sequencing in aquaculture breeding and genetics. Rev. Aquac. 1–13.

Rojas, V., N. Galanti, N. C. Bols, and S. H. Marshall, 2009 Productive infection of *Piscirickettsia salmonis* in macrophages and monocyte-like cells from rainbow trout, a possible survival strategy. J. Cell. Biochem. 108: 631–637.

Rozas, M., and R. Enríquez, 2014 Piscirickettsiosis and *Piscirickettsia salmonis* in fish: a review. J. Fish Dis. 37: 163–188.

Sernapesca, 2016a Informe Sanitario de Salmonicultura en Centros Marinos 2015. Servicio Nacional de Pesca y Acuicultura.

Sernapesca, 2016b Informe sobre uso de antimicrobianos en la salmonicultura nacional.

Sonesson, A. K., and T. H. E. Meuwissen, 2009 Testing strategies for genomic selection in aquaculture breeding programs. Genet. Sel. Evol. 41: 37.

Stear, M. J., S. C. Bishop, B. a Mallard, and H. Raadsma, 2001 The sustainability, feasibility and desirability of breeding livestock for disease resistance. Res. Vet. Sci. 71: 1–7.

Svishcheva, G. R., T. I. Axenovich, N. M. Belonogova, C. M. van Duijn, and Y. S. Aulchenko, 2012 Rapid variance components–based method for whole-genome association analysis. Nat. Genet. 44: 1166–1170.

Thompson, E. A., and R. G. Shaw, 1990 Pedigree Analysis for Quantitative Traits: Variance Components without Matrix Inversion. Biometrics 46: 399–413.

Tsai, H.-Y., A. Hamilton, A. E. Tinch, D. R. Guy, J. E. Bron et al., 2016 Genomic prediction of host resistance to sea lice in farmed Atlantic salmon populations. Genet. Sel. Evol. 48: 47.

Tsai, H. Y., A. Hamilton, A. E. Tinch, D. R. Guy, K. Gharbi et al., 2015 Genome wide association and genomic prediction for growth traits in juvenile farmed Atlantic salmon using a high density SNP array. BMC Genomics 1–9.

Vallejo, R. L., T. D. Leeds, B. O. Fragomeni, G. Gao, A. G. Hernandez et al., 2016 Evaluation of genome-enabled selection for bacterial cold water disease resistance using progeny performance data in rainbow trout: Insights on genotyping methods and genomic prediction models. Front. Genet. 7: 1–13.

Vallejo, R. L., T. D. Leeds, G. Gao, J. E. Parsons, K. E. Martin et al., 2017 Genomic selection models double the accuracy of predicted breeding values for bacterial cold water disease resistance compared to a traditional pedigree-based model in rainbow trout aquaculture. Genet. Sel. Evol. 49: 17.

VanRaden, P. M., 2008 Efficient methods to compute genomic predictions. J. Dairy Sci. 91: 4414–23.

Yañez, J. M., R. Bangera, J. P. Lhorente, A. Barria, M. Oyarzun et al., 2016a Negative genetic correlation between resistance against Piscirickettsia salmonis and harvest weight in coho salmon (Oncorhynchus kisutch). Aquaculture 459: 8–13.

Yañez, J. M., S. Naswa, M. E. Lopez, L. Bassini, K. Correa et al., 2016b Genomewide single nucleotide polymorphism discovery in Atlantic salmon (Salmo salar): validation in wild and farmed American and European populations. Mol. Ecol. Resour. 16: 1002–1011.

Yáñez, J.M. S. Newman, and R. D. Houston, 2015 Genomics in aquaculture to better understand species biology and accelerate genetic progress. Front. Genet. 6: 1–3.

Yáñez, J. M., R. D. Houston, and S. Newman, 2014a Genetics and genomics of disease resistance in salmonid species. Front. Genet. 5: 1–13.

Yáñez, J. M., J. P. Lhorente, L. N. Bassini, M. Oyarzún, R. Neira et al., 2014b Genetic co-variation between resistance against both Caligus rogercresseyi and Piscirickettsia salmonis, and body weight in Atlantic salmon (Salmo salar).

Yáñez, J. M., L. N. Bassini, M. Filp, J. P. Lhorente, R. W. Ponzoni et al., 2014c Inbreeding and effective population size in a coho salmon (Oncorhynchus kisutch) breeding nucleus in Chile. Aquaculture 420–421: S15–S19. Aquaculture 433: 295–298.

Yáñez, J. M., R. Bangera, J. P. Lhorente, M. Oyarzún, and R. Neira, 2013 Quantitative genetic variation of resistance against Piscirickettsia salmonis in Atlantic salmon (Salmo salar). Aquaculture 414–415: 155–159.

Yañez, J. M., and V. Martinez, 2010 Genetic factors involved in resistance to infectious diseases in salmonids and their application in breeding programmes. Arch. Med. Vet. 42: 1–13.

